# Calprotectin-mediated zinc chelation inhibits *Pseudomonas aeruginosa* protease activity in cystic fibrosis sputum

**DOI:** 10.1101/2021.02.25.432981

**Authors:** Danielle M. Vermilyea, Alex W. Crocker, Alex H. Gifford, Deborah A. Hogan

## Abstract

*Pseudomonas aeruginosa* induces pathways indicative of low zinc availability in the cystic fibrosis (CF) lung environment. To learn more about *P. aeruginosa* zinc access in CF, we grew *P. aeruginosa* strain PAO1 directly in expectorated CF sputum. The *P. aeruginosa* Zur transcriptional repressor controls the response to low intracellular zinc, and we used the NanoString methodology to monitor levels of Zur-regulated transcripts including those encoding a zincophore system, a zinc importer, and paralogs of zinc containing proteins that do not require zinc for activity. Zur-controlled transcripts were induced in sputum-grown *P. aeruginosa* compared to control cultures, but not if the sputum was amended with zinc. Amendment of sputum with ferrous iron did not reduce expression of Zur-regulated genes. A reporter fusion to a Zur-regulated promoter had variable activity in *P. aeruginosa* grown in sputum from different donors, and this variation inversely correlated with sputum zinc concentrations. Recombinant human calprotectin (CP), a divalent-metal binding protein released by neutrophils, was sufficient to induce a zinc-starvation response in *P. aeruginosa* grown in laboratory medium or zinc-amended CF sputum indicating that CP is functional in the sputum environment. Zinc metalloproteases comprise a large fraction of secreted zinc-binding *P. aeruginosa* proteins. Here we show that recombinant CP inhibited both LasB-mediated casein degradation and LasA-mediated lysis of *Staphylococcus aureus*, which was reversible with added zinc. These studies reveal the potential for CP-mediated zinc chelation to post-translationally inhibit zinc metalloprotease activity and thereby impact the protease-dependent physiology and/or virulence of *P. aeruginosa* in the CF lung environment.

**Importance:** The factors that contribute to worse outcomes in individuals with cystic fibrosis (CF) with chronic *Pseudomonas aeruginosa* infections are not well understood. Therefore, there is a need to understand environmental factors within the CF airway that contribute to *P. aeruginosa* colonization and infection. We demonstrate that growing bacteria in CF sputum induces a zinc-starvation response that inversely correlates with sputum zinc levels. Additionally, both calprotectin and a chemical zinc chelator inhibit the proteolytic activities of LasA and LasB proteases suggesting that extracellular zinc chelators can influence proteolytic activity and thus *P. aeruginosa* virulence and nutrient acquisition *in vivo*.

## Introduction

In cystic fibrosis (CF), microbes such as *Pseudomonas aeruginosa* colonize airway mucus where they then compete with host cells and other microbes for nutrients, including metals. Divalent metal ions (e.g., Zn^2+^, Fe^2+^, Mn^2+^, etc.) are essential micronutrients for host and microbe alike, in part, because they act as cofactors in enzymes important for a variety of cellular functions. While the concentration of metals, such as zinc, in CF sputum can vary, the concentration of zinc in expectorated sputum from CF patients is elevated, on average, compared to levels in samples from healthy controls (1-3). However, studies investigating the transcriptional response of *P. aeruginosa* in CF sputum show that a common gene expression pattern is the increased expression of zinc uptake and transport genes (4-9), which are normally expressed when zinc is limited. The *P. aeruginosa* zinc-starvation response is regulated by the zinc uptake regulator (Zur), which is a transcriptional repressor (10). When intracellular zinc is high, Zur monomers bind zinc, dimerize, and bind DNA to repress gene expression of zinc uptake pathways. When intracellular zinc becomes low, the dimeric, zinc-bound fraction of Zur decreases, which leads to derepression of genes involved in zinc uptake and the expression of zinc-free paralogs of essential proteins (zinc-sparing response). The *P. aeruginosa* Zur regulon (11, 12) includes the zinc transporter-encoding operon *znuABCD* (10, 13, 14), the zincophore-encoding operon *cntILMO* (15, 16), and zinc-free paralogs of ribosomal proteins (*PA3600* and *PA3601*) (13, 17) and transcription factors (*dksA2*) (18). These responses not only reduce the requirement for zinc but liberate the zinc that was stored in the zinc-dependent forms of these proteins (19).

The host, on the other hand, utilizes nutritional immunity to sequester metal ions away from pathogens to reduce bacterial growth and control infection (20). One of the most abundant zinc-binding host proteins in CF is calprotectin (CP), which was previously named “the cystic fibrosis antigen” because of its abundance in the serum, sputum, and bronchoalveolar lavage fluid (BALF) of individuals with CF (2, 21-24). Neutrophils recruited to sites of inflammation release CP as S100A8/A9 heterodimers (25, 26), which then form tetramers in environments with sufficient levels of calcium (27, 28). Each heterodimer has two divalent-metal binding sites: one site has high affinity for zinc and low affinity for manganese while the other site is capable of binding divalent manganese, iron, zinc, or nickel (29). CP is thought to induce zinc limitation as a means to control infections caused by *Staphylococcus aureus, Acinetobacter baumannii* in tissues, and *Salmonella enterica* serovar Typhimurium in the gastrointestinal tract (30-32). However, little is known about the effect of CP-mediated zinc sequestration on *P. aeruginosa* growth and physiology.

Additionally, CP has been shown to inhibit the activity of metalloproteases such as host matrix metalloproteinases via zinc chelation (33). *P. aeruginosa* regulates expression of several metalloenzymes, including zinc metalloproteases, by quorum sensing (QS), which is a mechanism that regulates gene expression in accordance with cell density through the secretion of signal molecules. The secretion of zinc metalloproteases LasB (PA3724), LasA (PA1871), AprA (PA1249), ImpA (PA0572), PepB (PA2939), and Protease IV (PA4175) (**Table 1**) are regulated by transcriptional regulators LasR and RhlR involved in QS (34, 35). This coordinated expression may be of particular importance for optimal protease activity given recent findings showing that LasB, Protease IV, and LasA are activated after being secreted by a QS-induced proteolytic cascade in which LasB activates Protease IV and then Protease IV, in turn, activates LasA (36, 37). Expression of these zinc metalloproteases is important for *P. aeruginosa* colonization and virulence because they play key roles in processes such as degrading host proteins (e.g., elastin) (38), invading host cells (39), evading host immune responses (40-42), and lysing other bacteria (e.g., *S. aureus*) (43, 44). While incubation of *P. aeruginosa* zinc metalloproteases with chemical zinc chelators inhibits their activity (45, 46), the effect of physiologically relevant zinc chelators such as CP on the activity of *P. aeruginosa* zinc metalloproteases remains unclear.

**Table 1.** Zinc metalloproteases secreted by *P. aeruginosa*

| Gene Number<sup>a</sup> | PDB Entry<sup>b</sup> | Protein Name and Description |
| --- | --- | --- |
| PA0572 | 5KDW | ImpA, immunomodulating metalloprotease of *P. aeruginosa* |
| PA1249 | 1KAP | AprA, alkaline metalloprotease or aeruginolysin |
| PA1871 | 3IT5 | LasA, staphylolytic protease |
| PA2939 | N/A | PepB or PaAP, aminopeptidase |
| PA3724 | 1EZM | LasB, elastase or pseudolysin |
| PA4175 | N/A | Protease IV, endoprotease |

<sup>a</sup> From *P. aeruginosa* genome website, https://www.pseudomonas.com/.
<sup>b</sup> From Protein Data Bank (PDB) website, https://www.rcsb.org/.

To test these hypotheses, we used a novel method in which *P. aeruginosa* strain PAO1 was grown directly in unamended expectorated CF sputum and matched sputum samples treated with divalent metals (e.g., Zn^2+^ and Fe^2+^) and zinc chelators (e.g., TPEN and CP). The effect of zinc chelators on *P. aeruginosa* zinc metalloprotease activity was further assessed using protease-specific assays. Overall, our findings support a model in which zinc chelation by CP in the mucus of the CF lung may impact the ecology of colonizing *P. aeruginosa* by inhibiting the activity of proteases involved in processes such as nutrient acquisition and interspecies competition.

## Results

### *P. aeruginosa* exhibits a Zur-regulated zinc-starvation response when grown in CF sputum samples from different donors

Given that recent studies show that *P. aeruginosa* increases expression of Zur-regulated genes in CF sputum (4-6), we first constructed a *lacZ* fusion to the promoter of *PA3600* on the chromosome of *P. aeruginosa* strain PAO1 (PAO1 *att*::P*_PA3600_-lacZ*) to act as a tool to explore factors that influence the activation of the Zur regulon. *PA3600* encodes the Zur-regulated zinc-independent isoform of the 50s ribosomal protein L36 (11-13, 17). Activation of the *PA3600* promoter was first confirmed by measuring activity by *P. aeruginosa* grown in culture medium (LB), medium containing TPEN (*N,N,N’,N’*-tetrakis-2-pyridylmethyl-ethylenediamine), or medium containing both TPEN and zinc (**Fig. 1a**). TPEN is a membrane permeable metal ion chelator with a high affinity for zinc (47) and was therefore used to induce a zinc-starvation response in *P. aeruginosa. P. aeruginosa* grown for 3 h in LB had little promoter activity (~23 Miller Units [MU]), while growth in medium containing TPEN resulted in a seven-fold increase in promoter activity (~150 MU) (**Fig. 1a**). The addition of TPEN and an excess of zinc (1 mM) did not stimulate promoter activity (**Fig. 1a**). The ability of sputum to activate the *PA3600* promoter was then determined by growing *P. aeruginosa* in M63 minimal medium containing 0.2% glucose (M63), culture medium plus TPEN (positive control), or expectorated CF sputum from 10 different donors (**Fig. 1b**). While *P. aeruginosa* grown for 3 h in M63 exhibited greater promoter activity (~85 MU) than when grown in LB (~23 MU), growth in CF sputum resulted in a three-fold increase in promoter activity (~281 MU). Average promoter activation in CF sputum was statistically the same as promoter activity induced by TPEN (**Fig. 1b**).

**Fig. 1.**
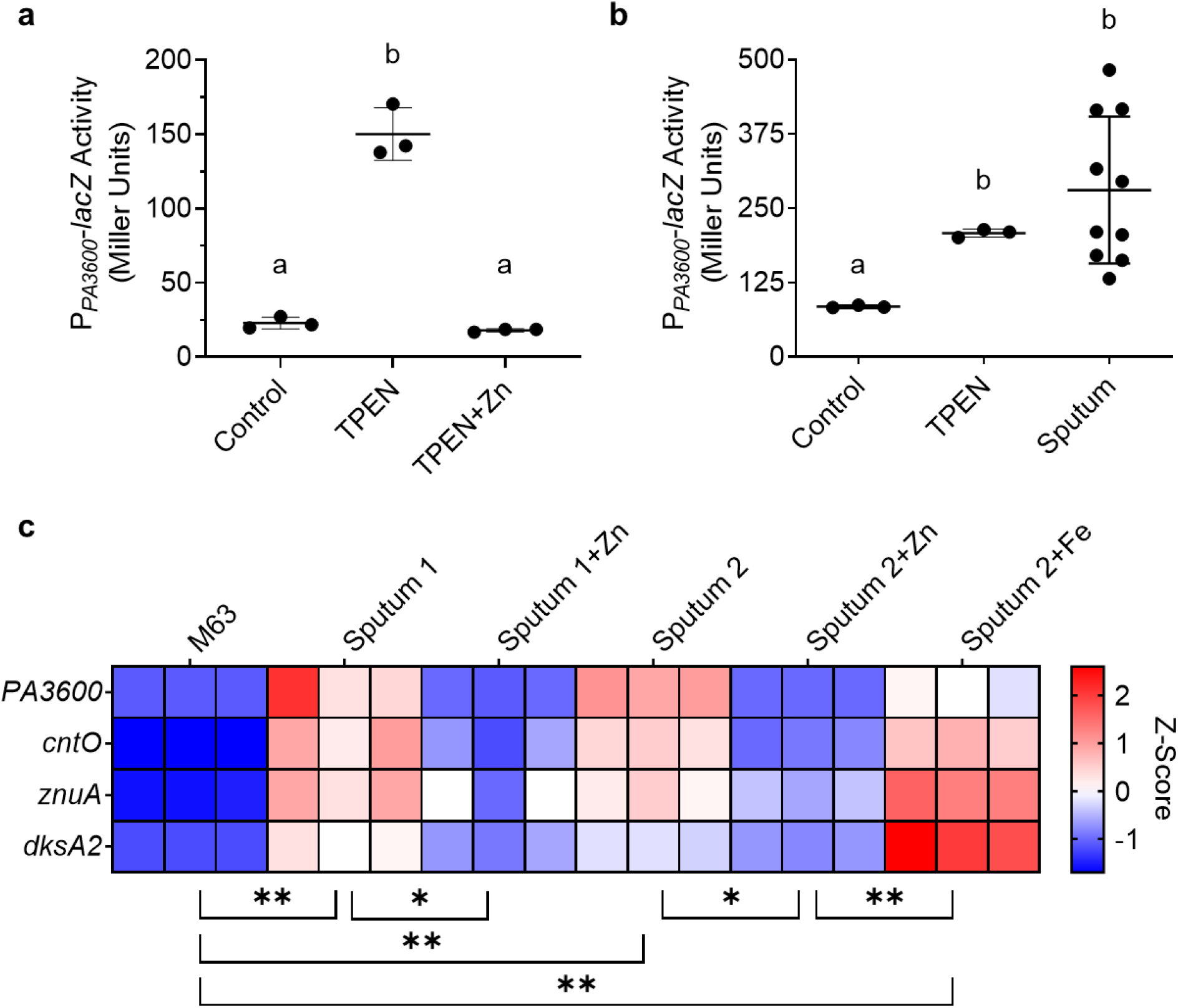
*P. aeruginosa* inoculated into expectorated CF sputum from different donors exhibits a zinc-starvation response. (**a**) *P. aeruginosa* strain PAO1 *P_PA3600_-lacZ* was grown in LB (Control), LB with 50 μM TPEN (TPEN), or LB with 50 μM TPEN and 1 mM ZnSO_4_ • 7 H_2_O (TPEN+Zn) for 3 h. The data shown represent the mean ± SD from three independent experiments. (**b**) *P. aeruginosa* strain PAO1 *P_PA3600_-lacZ* was grown in M63 (Control), M63 with 50 μM TPEN (TPEN), or expectorated CF sputum (sputum) for 3 h. Each point in the sputum set indicates a separate sample from a different donor. The data were analyzed by Brown-Forsythe and Welch ANOVA with Dunnett’s T3 multiple comparisons test. (**c**) *P. aeruginosa* strain PAO1 was inoculated into M63 (M63) or into sputum from two different donors (Sputum 1 and Sputum 2). The sputum was divided and left untreated (Sputum), treated with 1 mM ZnSO_4_ • 7 H_2_O (Sputum+Zn), or treated with 1 mM (NH_4_)_2_Fe(SO_4_)_2_ • 6 H_2_O (Sputum+Fe). Each condition was analyzed in triplicate. The same lowercase letters indicate samples that are not significantly different and different lowercase letters indicate significant differences (*p*<0.05). \**p*<0.05, \*\**p*<0.01

To further assess the activity of Zur in CF sputum, we used a multiplex method to assess expression of *PA3600* and three additional Zur-regulated genes. To do so, we used NanoString technology, which is a hybridization-based method that is quantitative, not hindered by contaminating DNA in sputum, and requires only a small amount of RNA. Consequently, NanoString works well for the analysis of small clinical sample aliquots (e.g., sputum) as previously demonstrated (48, 49). In this study, NanoString technology allowed for the analysis of subset of Zur-regulated genes: *PA3600, cntO, znuA,* and *dksA2*. Analysis showed an induction of these Zur-regulated genes in *P. aeruginosa* grown in sputum compared to M63 (**Fig. 1c**). Amending samples with excess zinc (1 mM) was sufficient to reduce the expression of Zur-regulated genes (**Fig. 1c**). Studies have shown regulatory crosstalk between iron and zinc as iron starvation was previously shown to increase expression of Zur-regulated genes *cntO, cntM*, and *amiA,* but not *znuA* (50). However, amending sputum samples with excess ferrous iron (1 mM) did not reduce expression of Zur-regulated genes (**Fig. 1c**). Together these data support the model that *P. aeruginosa* has limited access to zinc in sputum and that zinc and iron limitation are separate signals.

### Activation of the Zur-regulated *PA3600* promoter in CF sputum is inversely correlated with concentration of zinc in sputum samples

While promoter activity of *P. aeruginosa* grown in CF sputum samples was overall higher than medium controls, there was a range of promoter activity across sputum samples from different subjects (**Fig. 1b**). We hypothesized that differences in promoter activities between sputum samples from different CF patients were due to differences in sputum zinc concentrations. To test this, inductively coupled plasma mass spectrometry (ICP-MS) was performed on homogenized CF sputum samples to measure total metals (i.e., zinc, iron, and manganese) concentrations. The ability of these same sputum samples to activate the *PA3600* promoter in reporter strain PAO1 *att::P_PA3600_-lacZ* was tested in parallel. The data showed a significant inverse correlation between sputum zinc concentration and induction of the *PA3600* promoter across tested sputum samples (**Fig. 2**). There was no significant correlation between sputum iron or manganese concentrations and induction of the *PA3600* promoter (**Fig. 2b; Fig. S1a; Fig. S1b**). Induction of the *PA3600* promoter was also compared to clinical information, primarily lung function (FEV1%) at the time of sputum collection, but there was no correlation found between FEV1% and *PA3600* promoter activity (**Fig. S1c**). Therefore, the derepression of Zur-regulated genes in *P. aeruginosa* grown in CF sputum inversely correlates with the total zinc concentration in sputum samples.

**Fig. 2.**
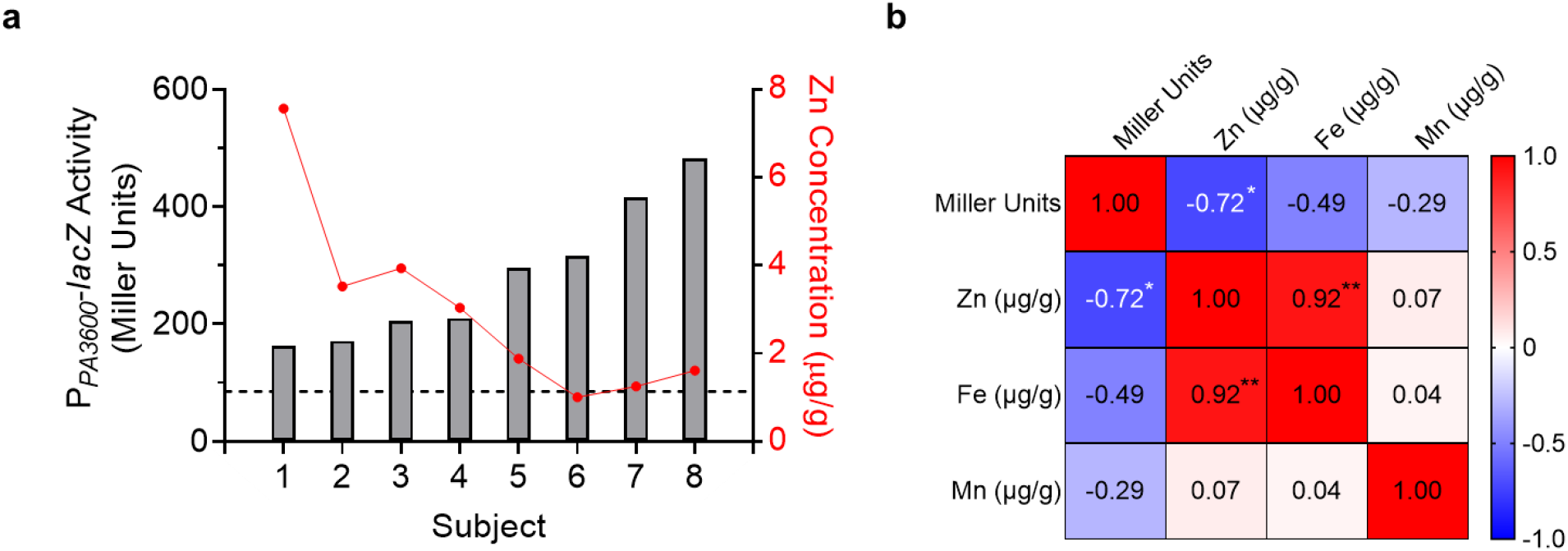
Activation of the *PA3600* promoter in CF sputum by *P. aeruginosa* is inversely correlated with total sputum zinc concentration. (**a**) *P. aeruginosa* strain PAO1 P*_PA3600_-lacZ* was inoculated into 8 different CF sputum samples. Zinc concentration of the same 8 CF sputum samples was determined by ICP-MS. B-Gal activity on the left y-axis (Miller Units; gray bars) was then compared to sputum zinc concentration on the right y-axis (μg/g; red dots), (**b**) Pearson correlation matrix comparing B-Gal activity (Miller units), sputum zinc concentration, sputum iron concentration, and sputum manganese concentration. \**p*<0.05, \*\**p*<0.01

### Recombinant CP induces a *P. aeruginosa* zinc-starvation response *in vitro* and in expectorated CF sputum

Studies report elevated levels of zinc in CF sputum (1, 2). Our ICP-MS data show that the sputum sample in our study that elicited the strongest zinc-starvation response had a zinc concentration of ~2 μg/g (~2000 μg/L, ~31 μM) (**Fig. 2a**). Given the concomitant high zinc concentration in our CF sputum samples and the elevated zinc starvation response in *P. aeruginosa* grown in these CF sputum samples, it is likely that the zinc in our CF sputum samples is bound by zinc-sequestering proteins. CP is one such host zinc-sequestering protein that is found in high concentrations in the sputum of CF patients (2, 22). CP has also been shown to induce expression of Zur-regulated genes in *P. aeruginosa* strain PA14 (51). Therefore, we hypothesized that CP binds zinc to induce a zinc starvation response in *P. aeruginosa* grown in CF sputum. To test this, we first expressed and purified recombinant human CP as previously described (52) and as illustrated in **Fig. S2**. The ability of our recombinant CP to induce a zinc-starvation response was tested by growing *P. aeruginosa* strain PAO1 *att::P_PA3600_-lacZ* in culture medium (LB), medium containing CP, or medium containing CP and zinc (**Fig. 3a**). CP concentrations in the lung can reach 1 mg/ml (~40 μM) (29), therefore, 1 mg/ml CP was used for all CP-based experiments. Growing *P. aeruginosa* in medium containing 1 mg/ml CP resulted in a four-fold increase in promoter activation (~92 MU) compared to the control (~25 MU) (**Fig. 3a**). The addition of excess zinc (1 mM) in the presence of CP prevented promoter activation (**Fig. 3a**). These results confirm that our purified recombinant human CP can induce a zinc-starvation response in *P. aeruginosa* which is quenched with the addition of exogenous zinc.

**Fig. 3.**
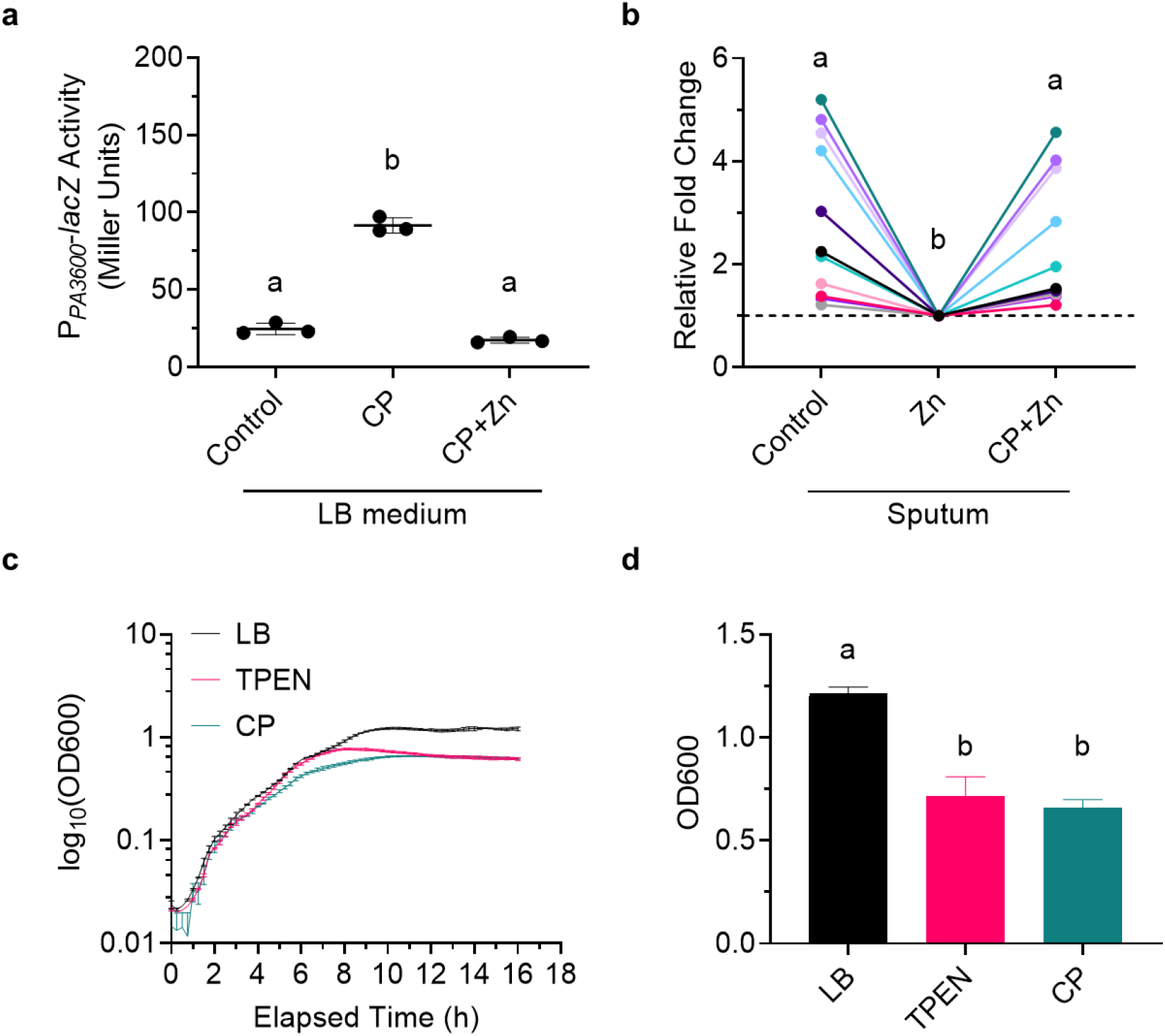
Recombinant human CP added to CF sputum and culture medium induces a zinc-starvation response by *P. aeruginosa*. (**a**) *P. aeruginosa* strain PAO1 P*_PA3600_-lacZ* was grown in culture medium (Control), medium with 40 μM CP (CP), or medium with 40 μM CP and 1 mM ZnSO_4_ • 7 H_2_O (CP+Zn) for 3 h. The data shown represent the mean ± SD from three independent experiments. (**b**) *P. aeruginosa* strain PAO1 *P_PA3600_-lacZ* was inoculated into CF sputum from 11 different donors. The sputum was divided and left untreated (Control), treated with 100 μM ZnSO_4_ • 7 H_2_O (Zn), or treated with 40 μM CP and 100 μM ZnSO_4_ • 7 H_2_O (CP+Zn) for 3 h. Different color dots represent samples from different donors. The same color dots connected by a line are from the same CF sputum donor. Data were analyzed by RM one-way ANOVA with Tukey’s multiple comparisons test. (**c**) Representative growth curves of *P. aeruginosa* strain PAO1 *P_PA3600_-lacZ* grown in LB, LB containing 50 μM TPEN, or LB containing 40 μM CP. Data shown represent the mean ± SD of three technical replicates and are representative of three independent experiments. (**d**) OD_600_ at 16 h of *P. aeruginosa* strain PAO1 *P_PA3600_-lacZ* grown in LB, LB containing 50 μM TPEN, or LB containing 40 μM CP. Data shown represent the mean ± SD of three independent experiments. The same lowercase letters indicate samples that are not significantly different and different lowercase letters indicate significant differences (*p*<0.05).

Despite the reportedly high concentrations of CP in the serum, sputum, and BALF of CF patients (2, 21-24), *P. aeruginosa* appears to be able to access enough zinc to persist. Various environmental factors may influence CP zinc binding such as calcium concentrations (53), pH (54), or the presence of oxidants (55, 56). Additionally, while CP in its tetrameric state is resistant to proteolytic degradation, CP is susceptible to oxidation which in turn makes it susceptible to proteolytic degradation by both host and bacterial proteases (55, 56). Because it was unclear if CP in sputum would remain intact and/or active to bind zinc, we tested the ability of recombinant human CP to bind zinc and thereby induce a zinc-starvation response in *P. aeruginosa* grown in CF sputum. *P. aeruginosa* strain PAO1 *att::P_PA3600_-lacZ* was grown in unamended CF sputum, sputum supplemented with 1 mM zinc, and sputum supplemented with both 1 mM zinc and 1 mg/ml (~40 μM) CP (**Fig. 3b**). The addition of zinc lowered *PA3600* promoter activity in sputum (**Fig. 3b**), supporting our NanoString data (**Fig. 1c**), while addition of CP to zinc-amended sputum significantly prevented reduction of promoter activity (**Fig. 3b**). These data confirm that recombinant CP added to CF sputum remains intact to bind zinc, which induces a zinc starvation response in colonizing *P. aeruginosa*.

While recombinant CP added to zinc-amended sputum increased *P. aeruginosa PA3600* promoter activity on average compared to zinc-amended sputum controls, the CF sputum samples tested varied in their responses (**Fig. 3b, Fig. 1b**). The inverse correlation between sputum zinc concentrations and induction of the *PA3600* promoter suggests that sputum samples that result in high promoter activity have lower concentrations of zinc than samples that induce low promoter activity, comparatively (**Fig. 2**). The high promoter activity by *P. aeruginosa* was readily quenched by the addition of zinc but remained high when CP was also added (**Fig. 3b**; green, lavender, lilac). Conversely, the low promoter activity by *P. aeruginosa* grown in sputum samples with presumably high zinc is not affected greatly by the addition of zinc nor CP (**Fig. 3b**; pink, light pink, gray). Overall, these data show that addition of recombinant CP to zinc-amended sputum can induce a zinc-starvation response dependent on sputum zinc concentration.

### Zinc metalloproteases are enriched amongst *P. aeruginosa-secreted* zinc-binding proteins

Since both TPEN and CP were confirmed to bind zinc and induce a zinc-starvation response in *P. aeruginosa* in culture medium (**Fig. 1a, Fig. 3a**), we wanted to further measure the effects of TPEN- and CP-mediated zinc sequestration on *P. aeruginosa* growth. Addition of TPEN or CP to cultures grown in LB decreased the final OD_600_ of *P. aeruginosa* compared to control conditions (**Fig. 3d**), but neither inhibited earlier growth stages (**Fig. 3c**). These data show that *P. aeruginosa* grows in the presence of CP under the conditions tested.

While CP does not prevent the growth of *P. aeruginosa*, little is known about how CP-mediated zinc starvation affects *P. aeruginosa* physiology. Unlike the chemical chelator TPEN, CP is not membrane permeable and instead exerts its effects on pathogens by binding metals in the extracellular environment. CF sputum has been reported to contain high concentrations of both CP (2, 22, 23) and secreted *P. aeruginosa* proteases including zinc metalloproteases (57). We performed a UniProt Knowledgebase (UniProtKB) analysis of the *P. aeruginosa* strain PAO1 proteome, which identified at least 72 zinc-binding proteins (**Table 2**). Of those 72, 64 were described by Gene Ontology (GO) molecular function as having catalytic activity (**Table 2**), which is consistent with the role of zinc as a cofactor. Of those 64 zinc-binding enzymes, 12 were further described as proteases and 5 of those were secreted zinc metalloproteases LasB, LasA, AprA, ImpA, and PepB (**Table 2**). We performed a second UniProtKB analysis of the *P. aeruginosa* strain PAO1 proteome that identified at least 34 secreted proteins, of which 6 were proteases and included the 5 aforementioned zinc metalloproteases in addition to Protease IV (PA4175). UniProtKB does not show Protease IV as binding zinc, but Protease IV has been described as a zinc metalloprotease and its enzymatic activity is reduced in a *P. aeruginosa* mutant lacking the zinc importer-encoding gene *znuA* (14). These analyses suggest that 83-100% of secreted proteases, important virulence factors, are zinc metalloproteases. Overall, previously published studies and curated databases suggest that CP and *P. aeruginosa*-secreted zinc metalloproteases are abundant in the extracellular milieu of the CF mucus environment.

**Table 2.** Characteristics of zinc-binding proteins in *P. aeruginosa* as annotated by UniProtKB^*a*^

| GO Molecular Function<sup>b</sup> | Number of Proteins | Subcellular Localization |  |  |  |
| --- | --- | --- | --- | --- | --- |
|  |  | Secreted | Inner Membrane | Cytoplasm | Not Listed |
| Zinc-Binding | 72 | 8 | 5 | 21 | 38 |
| Catalytic Activity: Non-peptidase | 52 | 3 | - | 16 | 33 |
| Catalytic Activity: Peptidase | 12 | 5 | 3 | 1 | 3 |
| Structural Binding Activity | 2 | - | - | - | 2 |
| Molecular Function Regulator | 2 | - | - | 1 | 1 |
| ATPase-Coupled Protein Transmembrane Transporter Activity | 1 | - | 1 | - | - |

<sup>a</sup> From protein knowledgebase (UniProtKB) website, https://www.uniprot.org/uniprot/.
<sup>b</sup> Gene Ontology (GO)

### Zinc chelation inhibits LasB-mediated proteolysis

Given the importance of zinc to the activity of zinc metalloenzymes, we hypothesized that zinc chelation by TPEN and CP would inhibit the activity of secreted zinc metalloproteases. Our initial studies suggested that LasB and LasA accounted for the majority of proteolytic activity by *P. aeruginosa* strain PAO1 (WT) because filtered supernatants from *ΔlasAB* cultures spotted onto milk plates cleared the milk plates substantially less than filtered WT supernatants (**Fig. 4a**, inset i-ii). As a result, this study focuses on the effect of zinc chelation on LasB and LasA activity.

**Fig. 4.**
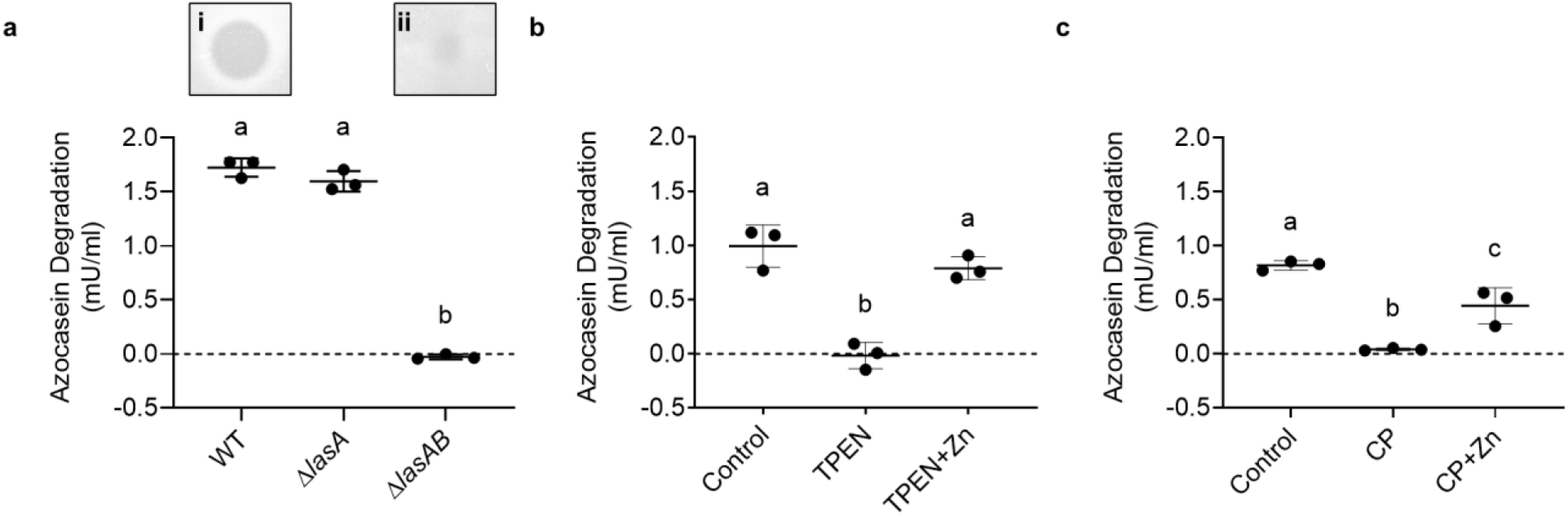
Zinc chelation inhibits LasB enzymatic activity. (**a**) Filtered supernatants from 16 h cultures of WT, *ΔlasA*, and *ΔlasAB* were incubated with 2% azocasein for 15 min. Inset are images showing the ability of (**i**) WT and (**ii**) *ΔlasAB* cell-free supernatants to clear milk plates after 16 h. (**b**) Filtered supernatants from WT 16 h cultures were left untreated (Control), treated with 50 μM TPEN (TPEN), or treated with 50 μM TPEN and 1 mM ZnSO_4_ • 7 H_2_O (TPEN+Zn) for an additional 16 h. Supernatants were then incubated with 2% azocasein for 15 min. The data shown represent the mean ± SD from three independent experiments. (**c**) Filtered supernatants from WT 16 h cultures were left untreated (Control), treated with 40 μM CP (CP), or treated with 40 μM CP and 1 mM ZnSO_4_ • 7 H_2_O (CP+Zn) for an additional 16 h. Supernatants were then incubated with 2% azocasein for 15 min. The same lowercase letters indicate samples that are not significantly different and different lowercase letters indicate significant differences (*p*<0.05). An enzyme unit (U) is defined as 1 μmol min^-1^.

To test the above hypothesis, LasB activity was determined quantitatively using azocasein as a substrate. The azocasein degradation assay was previously described to measure total proteolytic activity (14). However, by comparing the ability of *P. aeruginosa* WT, *ΔlasA*, and *ΔlasAB* supernatants to degrade azocasein, we found that azocasein degradation was LasB-dependent under the conditions tested (**Fig. 4a**). As a result, we tested the effect of TPEN and CP on LasB activity using the azocasein degradation assay. *P. aeruginosa* supernatants were filtered and then left untreated, treated with TPEN or CP, or treated with both TPEN or CP and zinc. Treatment with TPEN or CP inhibited LasB enzymatic activity while addition of excess zinc (1 mM) in the presence of TPEN or CP restored LasB activity (**Fig. 4b, Fig. 4c**). Furthermore, treatment of *ΔlasAB* supernatants with TPEN (**Fig. S3a**) or CP (**Fig. S3b**) without or with the addition of excess zinc did not alter azocasein degradation. Therefore, treatment of *P. aeruginosa* cell-free supernatants with zinc chelators TPEN and CP inhibits LasB-mediated caseinolytic activity.

### Zinc chelation inhibits LasA-mediated lysis of *S. aureus*

LasA activity was determined by monitoring the decrease in absorbance at 595 nm of a heat-killed *S. aureus* suspension as previously described (14). Use of *P. aeruginosa* strain PAO1 (WT), *ΔlasA*, and *ΔlasA+lasA* (complemented mutant) supernatants confirmed that LasA is necessary for the lysis of *S. aureus* and that this assay measures LasA-mediated lysis of *S. aureus* under the conditions tested (**Fig. 5a-b**). This assay was then used to measure LasA activity in *P. aeruginosa* cell-free supernatants left untreated, treated with TPEN or CP, or treated with both TPEN or CP and zinc. Treatment of supernatants with TPEN or CP inhibited LasA activity while treatment with TPEN or CP in the presence of excess zinc (500 μM and 160 μM, respectively) restored LasA activity (**Fig. 5c-f**). Furthermore, treatment of *ΔlasA* supernatants with zinc, TPEN, or CP had no effect on lysis of *S. aureus*, confirming that treatment of supernatants did not have LasA-independent cytotoxic effects on *S. aureus* (**Fig. S5**). Therefore, treatment of *P. aeruginosa* cell-free supernatants with zinc chelators TPEN and CP inhibits LasA-mediated lysis of *S. aureus*.

**Fig. 5.**
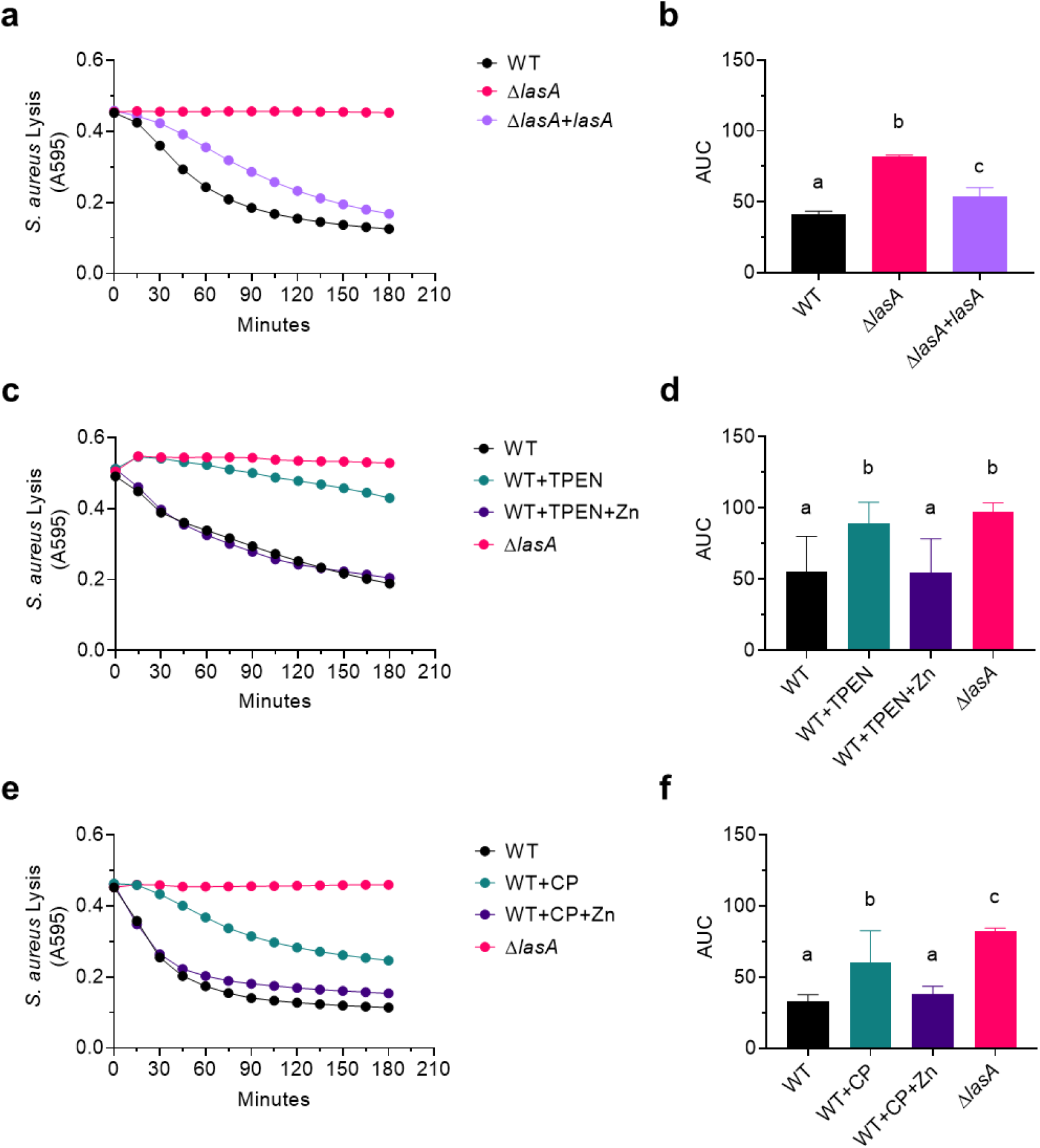
Zinc chelation inhibits LasA enzymatic activity. (**a-b**) Lysis of heat-killed *S. aureus* strain SH1000 by cell-free supernatants from WT, Δ*lasA*, and *ΔlasA+lasA (ΔlasA* complemented *in trans* under the Control of arabinose-inducible P_BAD_) 16 h cultures. (**c-d**) Lysis of heat-killed *S. aureus* strain SH1000 by WT and *ΔlasA* cell-free supernatants. WT supernatant was divided and left untreated (WT), treated with 50 μM TPEN (WT+TPEN), or treated with 50 μM TPEN and 500 μM ZnSO_4_ • 7 H_2_O (WT+TPEN+Zn). (**e-f**) Lysis of heat-killed *S. aureus* strain SH1000 by WT and Δ*lasA* cell-free supernatants. WT supernatant was divided and left untreated (WT), treated with 40 μM CP (WT+CP), or treated with 40 μM CP and 160 μM ZnSO_4_ • 7 H_2_O (WT+CP+Zn). (**a**), (**c**), (**e**) The data represent the mean from three independent experiments. Error bars have been omitted for clarity. (**b**), (**d**), (**f**) Quantification of data in (**a**), (**c**), and (**e**), respectively, using area under the curve (AUC). Data are the mean ± SD from three independent experiments. The same lowercase letters indicate samples that are not significantly different and different lowercase letters indicate significant differences (*p*<0.05).

## Discussion

Here we show that *P. aeruginosa* strain PAO1 grown in aliquots of expectorated CF sputum exhibits a zinc-starvation response despite relatively high concentrations of zinc in the sputum samples. Treatment with recombinant host CP was sufficient to induce a zinc-starvation response in *P. aeruginosa* grown in zinc-amended CF sputum samples from different subjects, demonstrating that CP retains its function in sputum. Furthermore, treatment of *P. aeruginosa* supernatants with CP inhibited the activity of secreted, extracellular zinc metalloproteases LasB and LasA. The data presented in this study support a model in which CP released from recruited neutrophils sequesters zinc from the environment to induce a zinc-starvation response in *P. aeruginosa* and sequesters zinc from secreted virulence factors including zinc-dependent metalloproteases LasA and LasB inhibiting *S. aureus* lysis, degradation of peptides, and/or nutrient acquisition (**Fig. 6**).

**Fig. 6.**
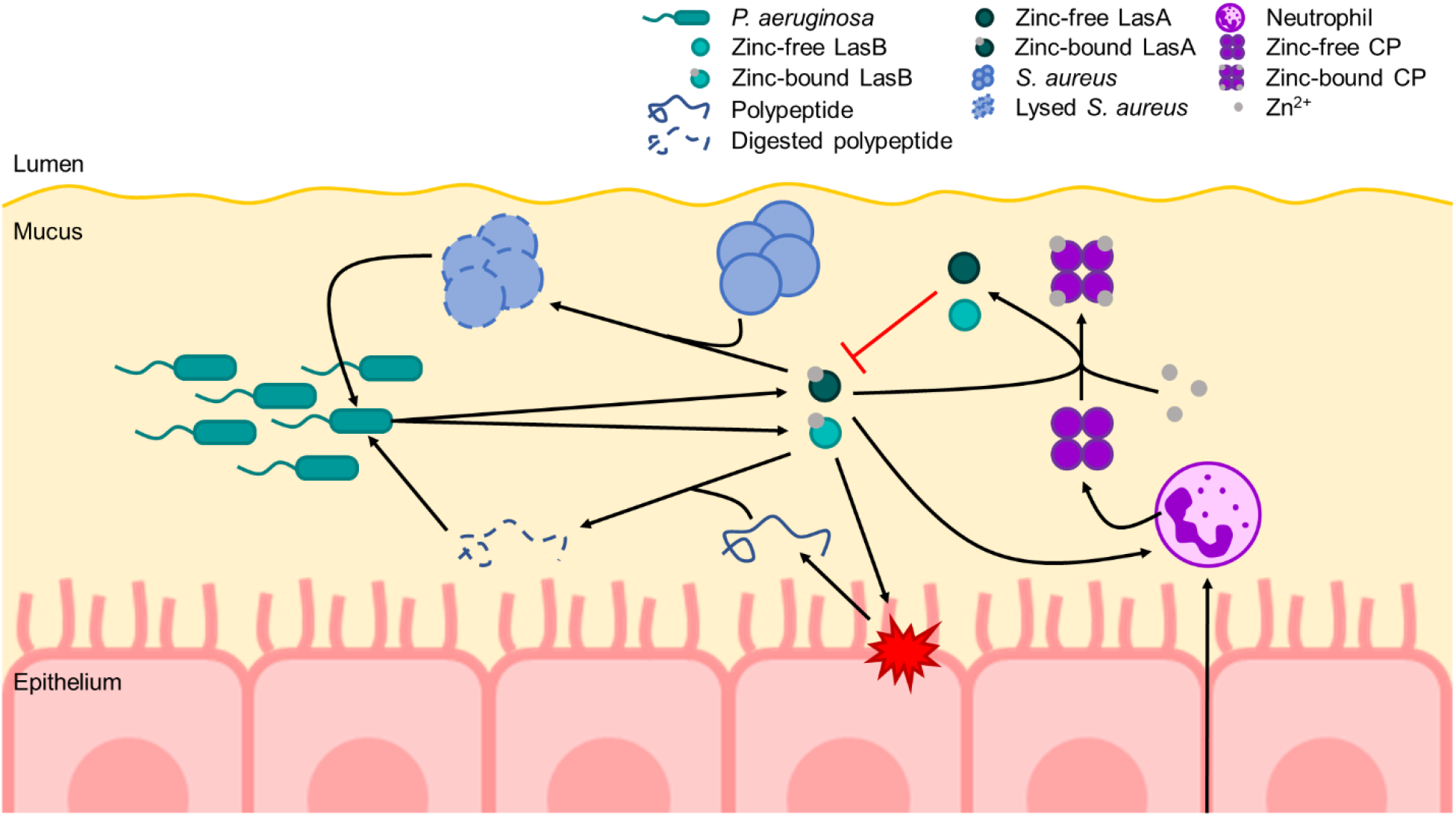
Model of the effects of CP-mediated zinc chelation in the CF lung on *P. aeruginosa. P. aeruginosa* colonizes the mucus in the airways of CF patients to high densities, which in part requires the uptake and utilization of zinc. At high densities, *P. aeruginosa* secretes a variety of quorum sensing-dependent virulence factors including zinc metalloproteases such as LasB and LasA. LasB is a protease that can degrade host proteins, such as elastin, as well as peptides. These degraded proteins/peptides can then be taken up and utilized as nutrients by *P. aeruginosa*. LasA is a protease that lyses *S. aureus* by cleaving pentaglycine bridges of peptidoglycan. LasA-mediated lysis of *S. aureus* allows *P. aeruginosa* to take up nutrients released from lysed *S. aureus* as well as to outcompete *S. aureus* in the CF lung. During infection, neutrophils are recruited to sites of infection/inflammation. Neutrophils may then release cellular contents such as CP. CP can then bind bioavailable zinc away from *P. aeruginosa* thus reducing the overall abundance of *P. aeruginosa*, while also inducing a zinc-starvation response by *P. aeruginosa*. Additionally, CP can bind zinc away from both LasB and LasA thereby inhibiting their proteolytic activity. Furthermore, LasB and LasA activity have been shown to induce neutrophil extracellular traps (NETs). Therefore, CP-mediated inhibition of LasB and LasA activity may lead to less NET formation and, subsequently, less CP release. Black arrows indicate a positive interaction. Red arrows indicate an inhibitory interaction.

A variety of strategies have been used to learn about the environment that *P. aeruginosa* encounters in the CF lung including analysis of bacteria grown in buffered media supplemented with CF sputum compared to bacteria grown in laboratory media (8, 9), and direct analysis of gene expression by bacteria in expectorated CF sputum (4, 5, 58). While studies have varied in their techniques, transcriptomic analyses have found that genes induced by low intracellular zinc are elevated in sputum samples relative to controls (4-9). Our model differs from previous models as it measures the transcriptional response of *P. aeruginosa* grown directly in expectorated sputum from a variety of CF patients. Our study also found that *P. aeruginosa* activates its zinc-starvation response in CF sputum on average but revealed differences across samples from different CF donors (**Fig. 1b-c, Fig. 3b**). These findings taken together underscore the fact that *P. aeruginosa* growth in laboratory media would not recapitulate the effect of low-zinc conditions in the context of CF. To this end, our CF sputum model is one way to provide a low-zinc environment and allows for investigation of the response of *P. aeruginosa* across sputum samples from different donors which vary in levels of host factors like CP. This same approach would also enable the investigation of different *P. aeruginosa* strains in sputum aliquots from a single donor.

CP concentrations during infections can reach 1 mg/ml or ~40 μM which is often posited to be higher than or in excess of the bioavailable zinc concentration in most environments (29). However, zinc concentrations in CF sputum are high relative to sputum from non-CF individuals and other biological compartments. Smith et al. (1) found that the zinc concentration of 45 CF sputum samples ranged from 678 μg/L (~10 μM) to 1181 μg/L (~18 μM) compared to 103 μg/L (~2 μM) to 597 μg/L (~9 μM) in 8 non-CF sputum samples. Li et al. (3) reported that the zinc concentration of 118 CF sputum samples ranged from ~5 μM to ~145 μM. In this study, the zinc concentration of 8 CF sputum samples ranged from 1.002 μg/g (~15 μM) to 7.562 μg/g (~116 μM) (**Fig. 2a**). Therefore, under certain conditions or in some microenvironments, CP may not be in excess of environmental zinc.

There is mounting evidence that divalent-metal sequestration by CP affects *P. aeruginosa*. Wakeman et al. (51) demonstrated that CP-mediated genetic responses in *P. aeruginosa* were reversed upon treatment with zinc *in vitro* and that *P. aeruginosa* and CP colocalized at sites of inflammation within a CF lung explant. D’Orazio et al. showed that CP-mediated growth inhibition was enhanced in *P. aeruginosa* strain Δ*znuA*, which is a mutant lacking the gene encoding the small zinc-binding protein of the ZnuABC zinc importer resulting in reduced intracellular zinc accumulation (13, 14). Zygiel et al. (59) showed that treatment with CP significantly reduced intracellular iron and manganese in *P. aeruginosa*, but did not significantly affect intracellular zinc, though intracellular zinc trended downward (59). Our data show that CP induces a Zur-regulated zinc-starvation response *in vitro* and in expectorated CF sputum which is repressed upon the addition of excess zinc (**Fig. 3a-b**). We also observed CP-mediated growth defects *in vitro* (**Fig. 3c**) similar to those reported by Zygiel et al. (59) which were previously attributed to ferrous iron chelation by CP. Taken together, the data show that *P. aeruginosa* and CP colocalize at sites of inflammation in the CF lung and that CP is capable of inducing zinc- and/or iron-starvation responses depending on test conditions.

Additionally, while Filkins et al. (60) showed that *in vitro* co-culture of *P. aeruginosa* and *S. aureus* on CF bronchial epithelial cells reduced the viability of *S. aureus*, Wakeman et al. (51) showed that zinc chelation by CP promotes *P. aeruginosa* and *S. aureus* co-culture in *in vitro, in vivo*, and *ex vivo* models, in part, by downregulating genes encoding anti-staphylococcal factors such as pyocyanin, hydrogen cyanide, and PQS/HQNO. Interestingly, treatment of *P. aeruginosa* with CP did not reduce the expression of *lasA* though the functionality of LasA was not tested (51). In this study, we show that CP-mediated zinc chelation inhibits LasA-mediated lysis of *S. aureus* by *P. aeruginosa in vitro* (**Fig. 5e-f**). Therefore, while LasA may be expressed and secreted by *P. aeruginosa* in the presence of CP, CP may post-translationally inhibit LasA activity via zinc sequestration. Furthermore, colonization of the CF airways is usually described as a pattern of succession where *S. aureus* is the predominant colonizer early on in younger patients before being outcompeted by *P. aeruginosa* in older patients (60). However, Fischer et al. (61) recently showed that *P. aeruginosa* and *S. aureus* chronically co-colonize the CF lung. Wakeman et al. also showed that *P. aeruginosa, S. aureus*, and CP colocalize in CF lung explants (51). Further studies are required to determine if CP modulates protease-dependent and/or protease-independent co-colonization of *P. aeruginosa* and *S. aureus* in the CF lung.

Notably, *P. aeruginosa* strains chronically adapted to the CF lung, including *lasR* loss-of-function (LasR-) mutants, have a reduced capacity to outcompete *S. aureus* (62). LasR is a QS regulator that positively regulates the expression and secretion of several virulence factors including zinc metalloproteases LasB, LasA, AprA, ImpA, PepB, and Protease IV (34, 35). However, LasR-strains commonly arise during chronic CF infection and are associated with worse lung function (63-68). While LasR-strains are common in CF infections, virulence factors regulated by LasR such as zinc metalloproteases are still reported to be abundant in CF sputum (57). Recent work by Mould et al. showed that when LasR+ and LasR-strains were cocultured, the LasR+ strain increased production of RhlR-controlled virulence factors by the LasR-strain (69). Interestingly, LasB and LasA are reportedly regulated by both the LasR and RhlR QS regulators (35). Therefore, further investigation is needed to understand how intra- and interspecies interactions within populations colonizing the CF airway impact the secretion and function of virulence factors such as zinc metalloproteases LasB and LasA.

LasB is an abundant protease with broad substrate specificity that is implicated in amino acid liberation and consumption (70). In addition to nutrient acquisition, LasB also plays a role in the ability of *P. aeruginosa* to invade host epithelial cells (39) and to evade host immune responses via processes such as degrading cytokines (40). Interestingly, degradation of pro-inflammatory cytokines IL-8 and IL-6 by LasB reduces neutrophil recruitment and the overall IL-8 and IL-6 response (40). While LasB-mediated cytokine degradation has been reported to reduce neutrophil recruitment, LasB can also induce neutrophil extracellular traps (NETs) (71, 72). Neutrophils recruited to sites of inflammation can release CP through processes such as NET formation (73) and in this study we show that CP-mediated zinc chelation inhibits the activity of secreted LasB (**Fig. 4c**). Taken together, there appears to be a complex interplay between LasB, neutrophils, and CP during the course of infection which may contribute to exacerbations in CF. Furthermore, recent work suggests that secreted LasB activates Protease IV which then predominantly processes and activates LasA (36, 37). Therefore, CP-mediated inhibition of secreted LasB activity may have downstream effects on the processing and activity of other secreted zinc metalloproteases.

In conclusion, the results of our study show that CP can induce a zinc-starvation response in *P. aeruginosa* in CF sputum as well as chelate zinc to inhibit the activity of virulence-associated zinc metalloproteases. Future studies will focus on how competition for zinc in a zinc-limited or zinc-chelating environment such as CF mucus shapes polymicrobial infections and patient outcomes, particularly considering the observed variability in zinc concentration and availability across CF patients.

## Materials and Methods

### Strains and growth conditions

Bacterial strains and plasmids used in this study are listed in **Table S1**. *P. aeruginosa* and *Escherichia coli* strains were maintained on lysogeny broth (LB) (1% tryptone, 0.5% yeast extract, 0.5% NaCl) with 1.5% agar and routinely grown in LB on a roller drum at 37°C LB. *P. aeruginosa* plasmid strains were maintained by supplementing media with 300 μg/ml carbenicillin or 60 μg/ml gentamicin. *E. coli* plasmid strains were maintained by supplementing media with 100 μg/ml carbenicillin. *S. aureus* SH1000 was maintained on trypticase soy with 1.5% agar (TSA) or grown in trypticase soy broth (TSB) on a roller drum at 37°C. *Saccharomyces cerevisiae* strains for cloning were maintained on yeast-peptone-dextrose (YPD) medium with 2% agar.

### Construction of plasmids

Primers used for plasmid construction are listed in **Table S2**. All plasmids were sequenced at the Molecular Biology Core at the Geisel School of Medicine at Dartmouth. Plasmid GH121_P*_PA3600_*-*lacZ* (DH3229) was constructed using a *S. cerevisiae* recombination technique as previously described (74). Plasmid GH121_P*_pqsA_*-*lacZ* served as the vector backbone for this construct. GH121_P*_PA3600_-lacZ* was purified from yeast using Zymoprep™ Yeast Plasmid Miniprep II according to manufacturer’s protocol and transformed into electrocompetent *E. coli* strain S17 by electroporation. The plasmid was introduced into *P. aeruginosa* by conjugation and recombinants were obtained using sucrose counter-selection and genotype screening by PCR.

Complementation plasmid pMQ70_*lasA* was generated using the NEBuilder HiFi DNA assembly cloning kit (New England BioLabs). *P. aeruginosa* strain PAO1V *ΔlasA* was complemented *in trans* by inserting a functional copy of *lasA* amplified from PAO1V genomic DNA into plasmid pMQ70 under the control of the arabinose-inducible *BAD* promoter generating plasmid pMQ70_*lasA*. Plasmid pMQ70_*lasA* was transformed into *ΔlasA* by electroporation.

### Cystic Fibrosis (CF) sputum collection

Sputum samples were collected in accordance with protocols approved by the Committee for the Protection of Human Subjects at Dartmouth. Expectorated sputum samples used in this study were collected from adult subjects with CF during a routine office visit or upon admission for treatment of a disease exacerbation. Sputum samples were frozen upon collection and stored at −80°C until use.

### Beta-galactosidase (β-Gal) assay

*P. aeruginosa* cells with a promoter fusion to *lacZ* integrated at the *att* locus were grown in 5 mL cultures of LB at 37°C for 16 h. Overnight cultures were diluted 1:50 in 50 ml culture medium (LB or M63) and then grown to an OD_600_ of 0.5. The cells were then centrifuged at 4,500 x g for 10 min, resuspended in culture medium, centrifuged at 10,000 x g for 2 min, and then resuspended in 500 μl culture medium. Ten μl of cell suspension were added per 100 μl culture medium or sputum sample in a 2 ml microcentrifuge tube. Samples were incubated at 37°C with shaking for 3 h. β-Gal activity was measured as described by Miller (75) using 50 μl of sample.

### RNA isolation and NanoString analysis

Unamended sputum or sputum amended with 1 mM ZnSO_4_ • 7 H_2_O or (NH_4_)_2_Fe(SO_4_)_2_ • 6 H_2_O (100 μL) was added to 2 ml microcentrifuge tubes. *P. aeruginosa* strain PAO1 was grown in 5 mL cultures of LB at 37°C for 16 h. Overnight cultures were diluted 1:50 in 50 ml M63 minimal medium with 0.2% glucose and then grown to an OD_600_ of 0.5. The cells were then centrifuged at 4,500 x g for 10 min, washed with water, centrifuged, and then resuspended in 500 μl water. Ten μl of cell suspension were added per 100 μl M63 minimal medium with 0.2% glucose (control) or sputum sample in a 2 ml microcentrifuge tube. Samples were then incubated at 37°C with shaking for 3 h. TriZol (900 μl) was added to 100 μl sputum containing 10 μl of PAO1 cell suspension. Samples were stored overnight. RNA was prepared following DirectZol kit instructions and eluted in 50 μl water.

For NanoString, 5 μl of a 1:10 dilution of RNA was used. Diluted RNA was applied to the codeset PaV4 and processed as previously reported (49). Counts were normalized to the geometric mean of spiked-in technical controls. Normalized counts were used for Z-score calculations and heatmap construction.

### Measurement of zinc in sputum samples

Sputum samples for zinc analysis were stored at −80°C until processed. Sputum zinc was quantified by inductively coupled plasma-mass spectrometry (ICP-MS) following nitric acid digestion of organic material according to the method of Heck et al. and is expressed as μg zinc per g of sputum (76). ICP-MS was performed by the Dartmouth Trace Element Analysis (TEA) Core.

### Expression and Purification of recombinant calprotectin (CP)

Plasmid S100A8/A9 was obtained from Futami et al. (52) and recombinant CP was expressed and purified as previously described with minor modification. Plasmid S100A8/A9 was first confirmed by Sanger sequencing and then transformed into *E. coli* T7 Express cells. Transformed T7 Express cells were then grown in LB containing 100 μg/ml carbenicillin at 37°C with shaking and induced at about an OD_600_ of 0.5 with 0.5 mM β-D-1-thiogalactopyranoside (IPTG) for 3 h. Cultures were centrifuged at 13,260 x g for 10 min at 4°C. Supernatant was discarded. Cell pellets were resuspended in 30 ml wash solution (150 mM NaCl), transferred to a 50 ml conical tube, and then centrifuged at 3,210 x g for 10 min at 4°C. Supernatant was discarded. Pellets were weighed and then stored at −20°C.

Cell pellets were resuspended in 85 mL lysis buffer (50 mM Tris-HCl pH 7.5, 50 mM NaCl, 5 mM MgCl_2_) supplemented with Benzonase-HC to control viscosity of the sample. Cells were then lysed using the microfluidizer with 3 passages at 18,000 psi. Final volume was about 100 ml. 15% polyethylenimine (PEI) was added dropwise to a final concentration of 0.7% to precipitate nucleic acids (about 5 ml). Samples were then centrifuged at 23,280 x g for 10 min at 4°C. Pellet containing intact cells and precipitated nucleic acids was discarded. NH_4_SO_4_ (61.27 g) was added slowly to clarified supernatant (about 115 ml) while stirring at 4°C until a saturation of 80%. The sample become gradually turbid. Sample was stirred for an additional 30 min after complete saturation. Sample was then centrifuged at 23,280 x g for 10 min at 4°C. Supernatant was discarded and the pellet was dissolved in about 30 ml solubilization buffer (50 mM Tris-HCl pH 7.5, 30 mM dithiothreitol [DTT]) and incubated for 1 h at 37°C. Dissolved pellet was transferred to dialysis cassettes and dialyzed overnight in 50 mM sodium phosphate pH 6.0 at 4°C using 3.5 kDa cut-off dialysis cassettes to change buffer. Sample was then centrifuged at 23,280 x g for 10 min at 4°C to remove any pellet.

CP was then purified using a HiTrap SP column (stored in 20% ethanol). The column was washed with 5 column volumes (CV) of H_2_O at about 5 ml/min. The column was then washed with 5 CV of 100% SP Sepharose HP buffer B (50 mM sodium phosphate pH 6.0, 1 mM DTT, 1 M NaCl; filtered/degassed) at about 5 ml/min. The column was equilibrated with 10 CV of SP Sepharose HP buffer A (50 mM sodium phosphate pH 6.0, 1 mM DTT; filtered/degassed) at about 5 ml/min. A superloop was assembled with the appropriate volume for sample application. Sample was then loaded in the column using the superloop at 2.5 ml/min. The column was then washed with 10 CV of SP Sepharose HP buffer A at about 5 ml/min. The column was then washed with a step gradient of SP Sepharose HP buffer B: 5 CV of 5% SP Sepharose HP buffer B, 10 CV at 30% SP Sepharose HP buffer B and 5 CV at 100% SP Sepharose HP buffer B at about 5 ml/min.

Fractions were analyzed using SDS-PAGE (15% gel) and the appropriate fractions were then pooled.

CP was then purified using a HiLoad 26/600 Sephadex S75 and CP buffer (50 mM Tris-HCl pH 7.5, 150 mM NaCl, 1 mM DTT; filtered/degassed). Sample (about 13 ml) was loaded in a 50 ml superloop. Sample was then run on the HiLoad 26/600 Superdex 75p, a program composed of 2 CV equilibration, injection of 12 ml sample and elution with 1.2 CV at 2.6 ml/min. Flow rate is 2.6 ml/min and collection of 7 ml/tube. Tubes corresponding to three different fractions were pooled to make fractions F1_I, F2_I, and F3_I. All other tubes containing calprotectin from both HiTrap runs were concentrated using YM-10 Amicon centrifugal filters and re-loaded in the HiLoad 26/600 superdex 75 as before. Tubes corresponding to three different fractions were pooled to make fractions F1_II, F2_II, and F3_II. Samples from all six fractions were analyzed using SDS-PAGE (4-12% gel). Fractions F1_I and F1_II, F2_I and F2_II, and F3_I and F3_II were combined to make fractions F1, F2, and F3, respectively. Fractions were concentrated with YM-10 Amicon centrifugal filters. The final concentrations of the fractions were determined using a Bradford protein assay.

### Protease assays

*P. aeruginosa* culture supernatants were used for protease assays. 5 ml overnight cultures in LB were centrifuged at 4,500 x g for 10 min. Supernatants were then filter sterilized using a 0.22 μm syringe filter. For TPEN experiments, undiluted supernatants were used. For CP experiments, stored aliquots of CP were first diluted to 3 mg/ml in CP buffer without DTT (50 mM Tris-HCl pH 7.5, 150 mM NaCl). Then 1 part 3 mg/ml CP was added to 2 parts supernatant for a final concentration of 1 mg/ml.

Caseinolytic activity was determined qualitatively by spotting *P. aeruginosa* supernatants onto 1% milk plates or quantitatively using azocasein as a substrate as previously described (14) with modification. In brief, *P. aeruginosa* culture supernatants were treated overnight (16 h) with 50 μM TPEN or an equivalent volume of 100% EtOH, 1 mg/ml (~40 μM) CP or an equivalent volume of CP buffer without DTT, and/or 1 mM ZnSO_4_ • 7 H_2_O or an equivalent volume of diH_2_O. Treatment of WT supernatants with 50 μM to 2 mM ZnSO_4_ • 7 H_2_O was found not to affect LasB activity (**Fig. S3c**). The supernatants were then incubated at 37°C overnight (16 h). Supernatants (25 μl) were mixed with 150 μl 2% azocasein in 10 mM Tris-HCl, 8 mM CaCl_2_, pH 7.4. Samples were incubated at 37°C for 15 min. 228 μl of 10% TCA were added to each sample, vortexed, then incubated at room temperature for 15 min. Samples were then centrifuged for 10 min at 10,000 x g. Cleared supernatants (100 μl) were added to wells of a 96-well flat-bottom polystyrene plate containing 200 μl 1 M NaOH. Absorbance was read at 440 nm.

Staphylolytic activity was determined by monitoring the decrease in absorbance at 595 nm of a heat-killed *S. aureus* suspension as previously described (14) with modification. *S. aureus* strain SH1000 (77) was cultured in TSB overnight (16 h) at 37°C with rolling. Cultures were centrifuged at 4,500 x g for 10 min, resuspended in 20 mM Tris-HCl, pH 8.8 to a final OD_600_ of 1.0, and then killed by heating at 100°C for 30 min. Heat-killed *S. aureus* suspensions were cooled to room temperature before use. *P. aeruginosa* culture supernatants were treated overnight (16 h) with 50 μM TPEN or an equivalent volume of 100% EtOH, 1 mg/ml (~40 μM) CP or an equivalent volume of CP buffer without DTT, and/or 160-500 μM ZnSO_4_ • 7 H_2_O or an equivalent volume of diH_2_O. Because increasing concentrations of zinc were previously reported to inhibit LasA activity (46), an appropriate concentration of zinc to use in add-back experiments was determined experimentally. For undiluted WT supernatants, the addition of 500 μM zinc had no effect on LasA activity, while increasing concentrations of zinc inhibited LasA-mediated lysis of *S. aureus* (**Fig. S4a-b**). Therefore, we used 500 μM zinc for TPEN-based experiments. For CP-buffer diluted WT supernatants, the addition of 50 μM zinc had no effect on LasA activity, while increasing concentrations of zinc inhibited LasA-mediated lysis of *S. aureus* (**Fig. S4c-d**). However, a tetramer of CP can potentially bind up to four zinc ions. Therefore, to ensure that zinc would be in excess in CP-based experiments, we used 160 μM zinc which was four times the concentration of CP but still less than 250 μM zinc which was the concentration tested that started to inhibit LasA activity independent of CP. *P. aeruginosa* supernatants (20 μl) were added to 180 μl of heat-killed *S. aureus* in wells of a 96-well flat-bottom polystyrene plate. Staphylolytic activity was determined by monitoring the change in absorbance at 595 nm every 15 min for 3 h using a plate reader. The plate was shaken before each read.

### Statistical analysis

Statistical analysis was performed using GraphPad Prism 8 and results were expressed as the mean values plus or minus standard deviations. Unless otherwise noted, one-way analysis of variance (ANOVA) followed by Tukey’s multiple-comparison test was performed to determine statistical significance of the data. See the figure legends for other specific statistical tests used.

## Acknowledgements

We would like to thank Pat Occhipinti for generating the promoter-*lacZ* fusion reporter strain used in this study, Andreia Verissimo of the Institute for Biomolecular Targeting (bioMT) Molecular Tools Core (MTC) for her assistance in expressing and purifying recombinant calprotectin used in this study, and Nick Jacobs and Georgia Doing for their insightful comments and feedback during manuscript preparation. Research reported in this publication was supported by the Cystic Fibrosis Foundation (CFF) grants GIFFOR1610 and GIFFOR17Y5 awarded to A.H.G. and CFF grant HOGAN19G0 awarded to D.A.H. Support for the project was also provided by bioMT through NIH NIGMS grant P20 GM113132, the Dartmouth Cystic Fibrosis Research Center (DartCF) through NIH NIDDK grant P30 DK117469, the CFF Research Development Program (CF RDP) through CFF grant STANTO19R0, and the Dartmouth Trace Element Analysis (TEA) Core through NIH NIEHS P42 ES007373. The content of this publication is solely the responsibility of the authors and does not necessarily represent the official views of the funding sources.

## Notes

### Competing Interest Statement

The authors have declared no competing interest.

### Summary of Updates

Supplemental files updated

