## Supplemental Material for "Calprotectin-mediated zinc chelation inhibits *Pseudomonas aeruginosa* protease activity in cystic fibrosis sputum"

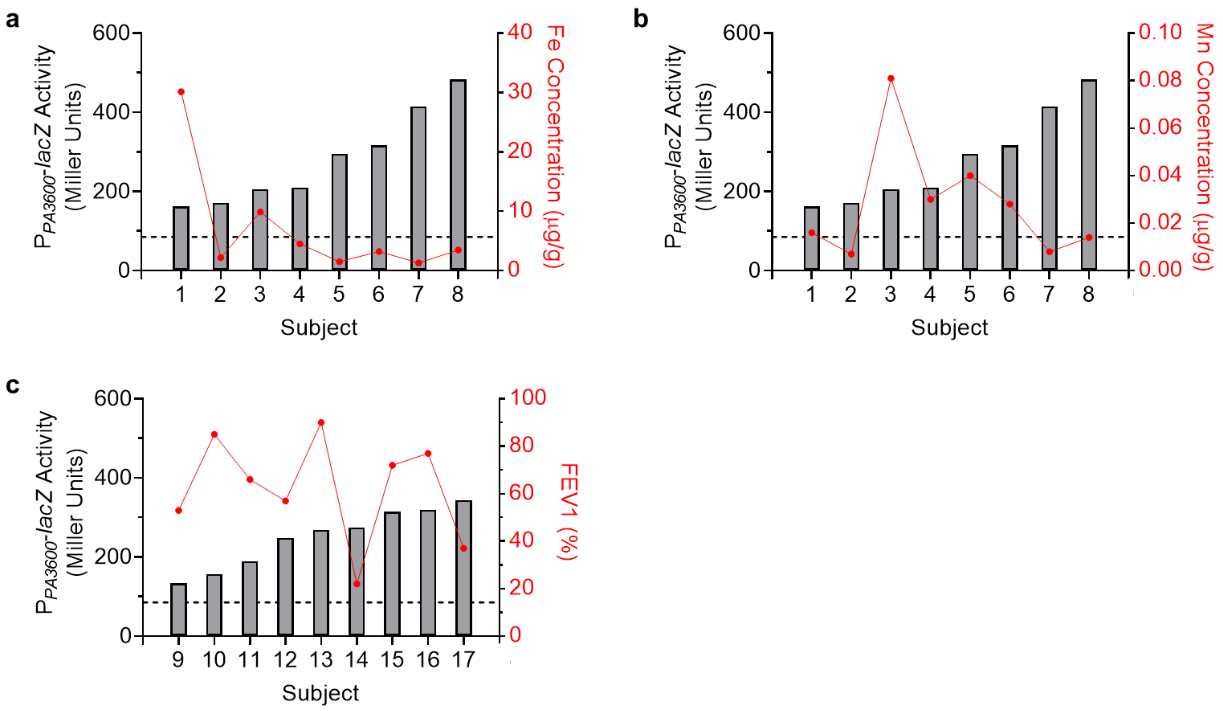


**Fig. S1** Activation of the *PA3600* promoter in CF sputum by *P. aeruginosa* is not correlated with total sputum iron or manganese concentration nor FEV1 of donors at the time of collection. *P. aeruginosa* strain PAO1 $P_{PA3600}$-*lacZ* was inoculated into 8 different CF sputum samples. (**a**) Iron and (**b**) manganese concentration of the same 8 CF sputum samples was determined by ICP-MS. B-Gal activity on the left y-axis (Miller units; gray bars) was then compared to sputum metal concentration on the right y-axis (µg/g; red dots). (**c**) *P. aeruginosa* strain PAO1 $P_{PA3600}$-*lacZ* was inoculated into an additional 9 different CF sputum samples. B-Gal activity on the left y-axis (Miller units; gray bars) was then compared to the FEV1 of the donors at the time of collection on the right y-axis (%; red dots).

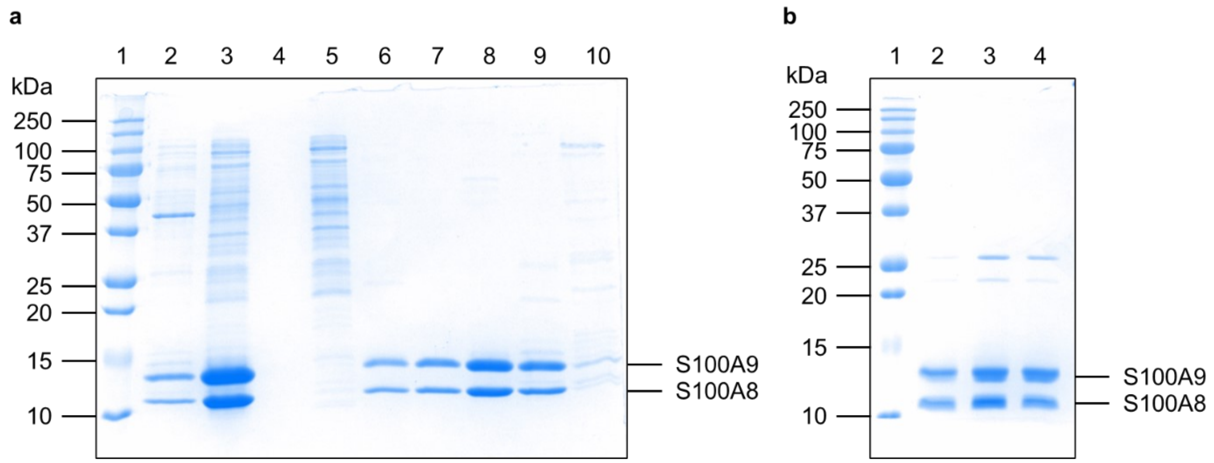


**Fig. S2** Expression, purification, and concentration of recombinant human CP. (**a**) SDS-PAGE using a 15% Laemmli gel performed on aliquots of the following samples collected during the process of purifying recombinant CP: Lane 1) Marker, Lane 2) Dialyzed solubilized pellet, Lane 3) Sample loaded on column, Lane 4) Empty, Lane 5) Tube 4, Lane 6) Tube 14, Lane 7) Tube 17, Lane 8) Tube 26, Lane 9) Tube 40, Lane 10) Tube 58. (**b**) SDS-PAGE using a 15% Laemmli gel performed on aliquots of pooled and concentrated CP-containing fractions. Lane 1) Marker, Lane 2) Fraction 1, Lane 3) Fraction 2, Lane 4) Fraction 3. Gels were stained with Coomassie Brilliant Blue.

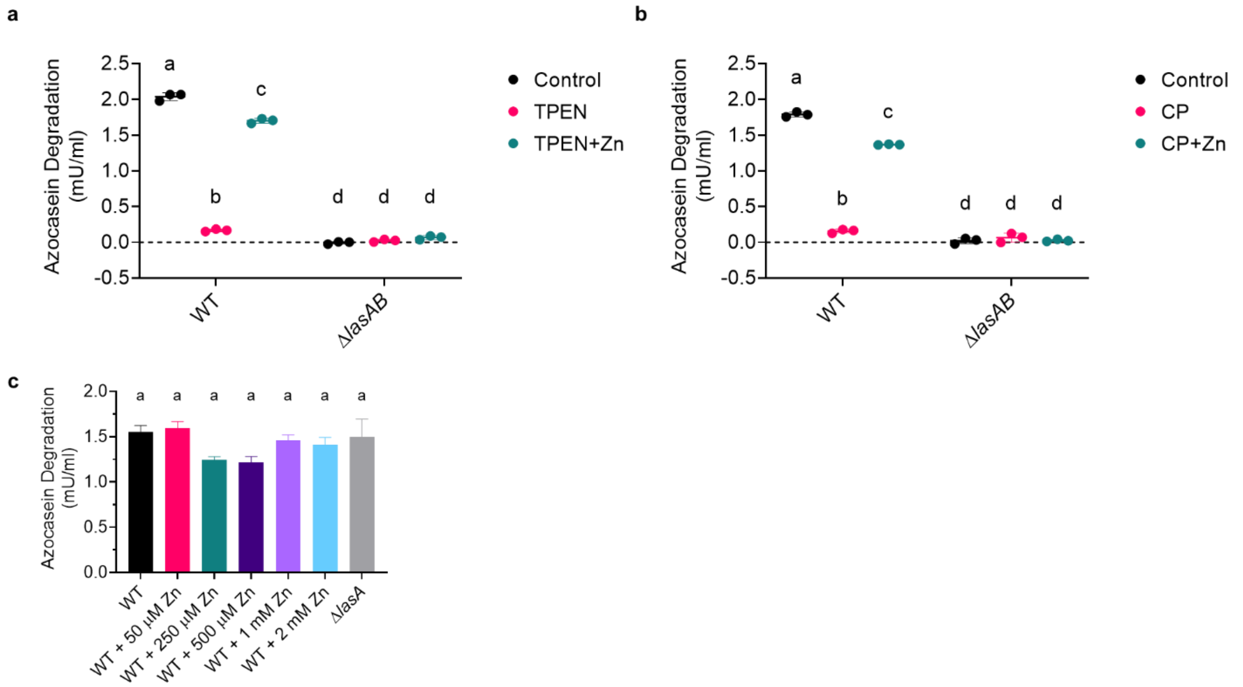


**Fig. S3** Treatment of Δ*lasAB* supernatants with zinc, TPEN, or CP does not degrade azocasein. Filtered supernatants from 16 h cultures of WT and Δ*lasAB* were left untreated (Control) or treated with (**a**) 50 μM TPEN (TPEN) or 50 μM TPEN plus 1 mM $ZnSO_4 \cdot 7 H_2O$ (TPEN+Zn) or (**b**) 40 μM CP or 40 μM CP plus 1 mM $ZnSO_4 \cdot 7 H_2O$ for an additional 16 h. Samples were then incubated with 2% azocasein for 15 min. The data shown represent the mean ± SD from three technical replicates from a representative experiment. Data were analyzed using two-way ANOVA with Sidak's multiple comparisons test. (**c**) Filtered supernatants from 16 h cultures of WT and Δ*lasA* were left untreated or treated with 50 μM to 2 mM $ZnSO_4 \cdot 7 H_2O$ for an additional 16 h. Samples were then incubated with 2% azocasein for 15 min. The data shown represent the mean ± SD from three independent experiments. The same lowercase letters indicate samples that are not significantly different and different lowercase letters indicate significant differences ($p<0.05$). An enzyme unit (U) is defined as 1 μmol $min^{-1}$.

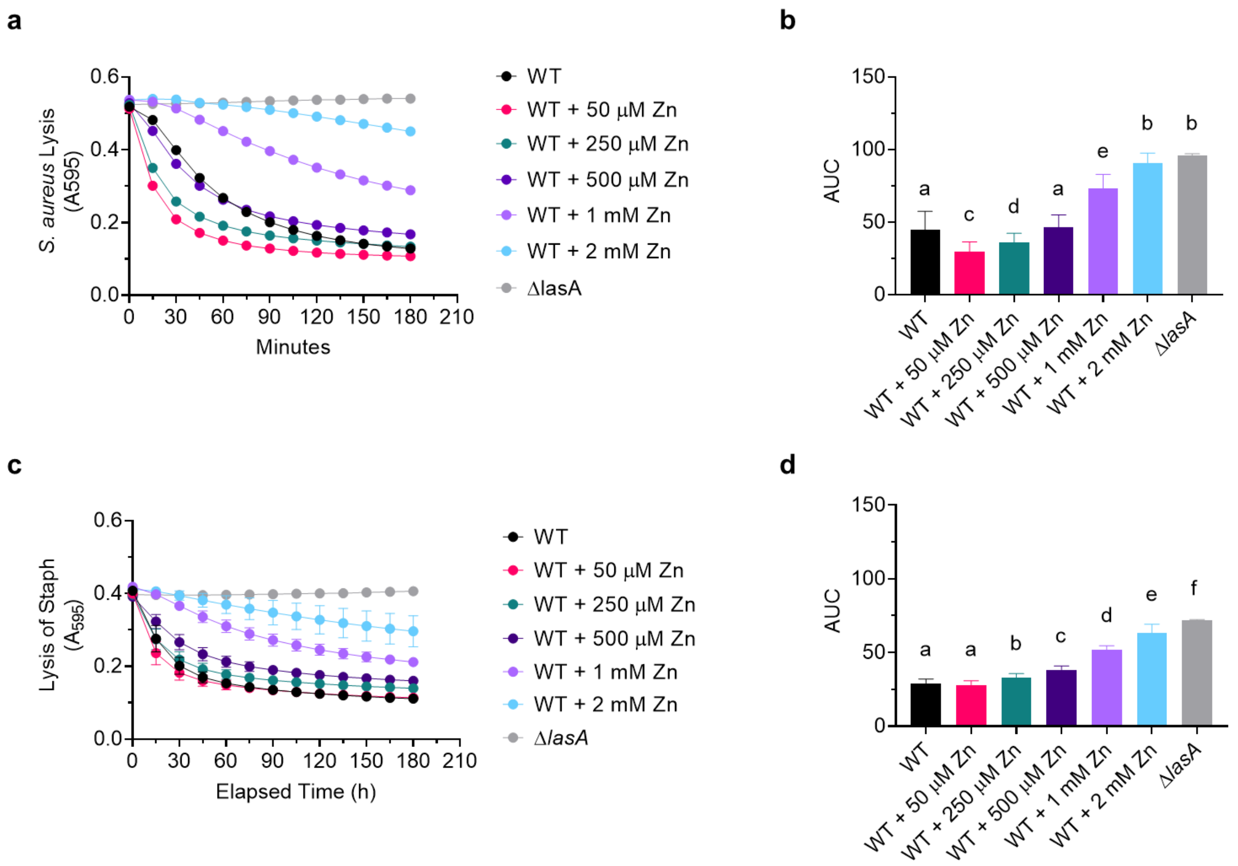


**Fig. S4** Treatment of *P. aeruginosa* filtered supernatants with increasing concentrations of zinc inhibits LasA activity. Lysis of heat-killed *S. aureus* strain SH1000 by WT and Δ*lasA* cell-free supernatants. WT supernatant was (**a**) left undiluted or (**b**) diluted two parts to one part CP buffer without DTT and then divided into aliquots and left untreated (WT) or treated with 50 μM to 2 mM $ZnSO_4 \cdot 7\ H_2O$. (**a**) and (**c**) The data represent the mean from three independent experiments. Error bars have been omitted for clarity. (**b**) and (**d**) Quantification of data in (**a**) and (**c**), respectively, using area under the curve (AUC). Data are the mean ± SD from three independent experiments. The same lowercase letters indicate samples that are not significantly different and different lowercase letters indicate significant differences ($p$<0.05).

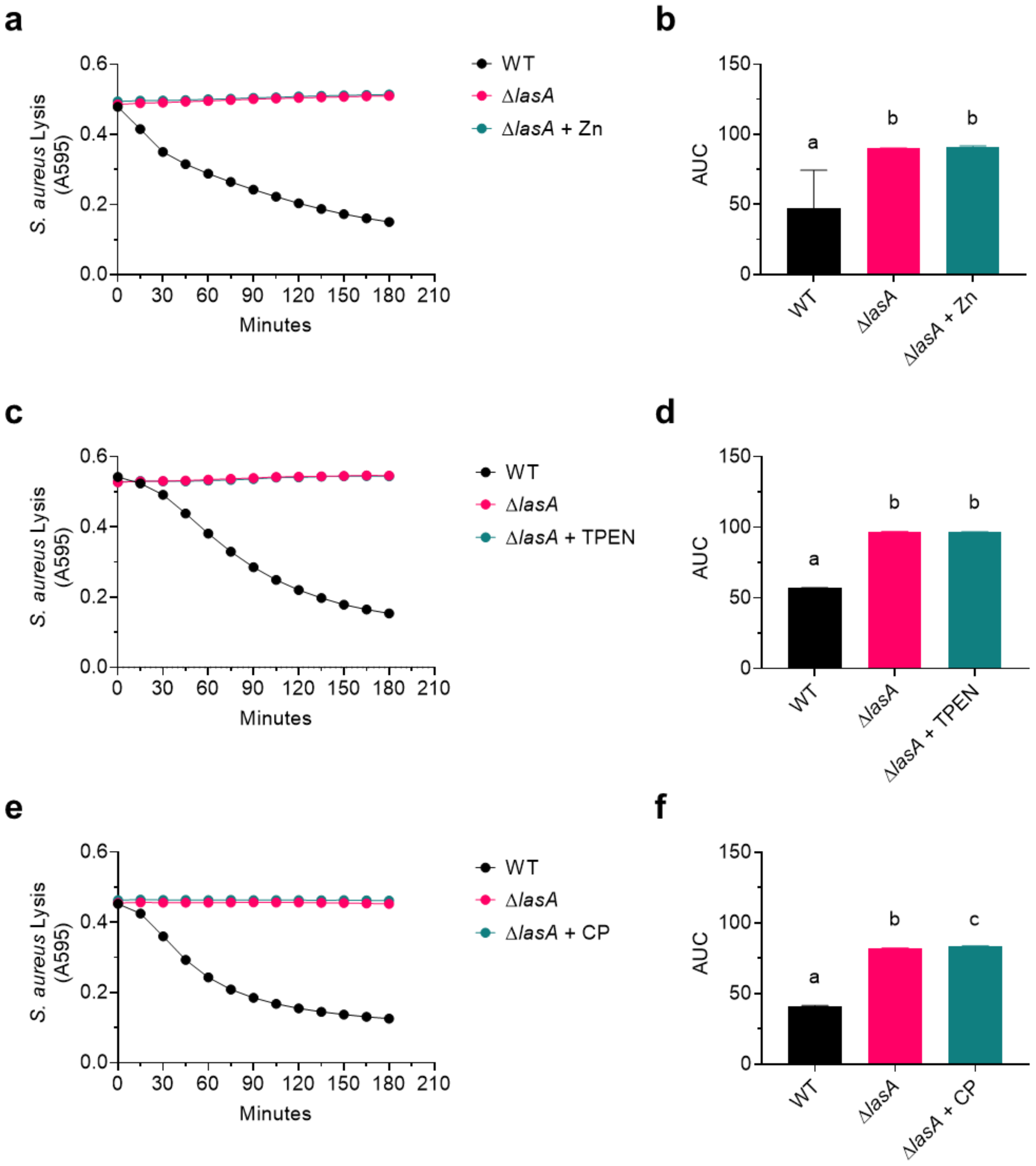


**Fig. S5** Δ*lasA* supernatants treated with zinc, TPEN, or CP do not lyse heat-killed *S. aureus*. Lysis of heat-killed *S. aureus* strain SH1000 by WT and Δ*lasA* cell-free supernatants treated with (**a**) 500 μM $ZnSO_4 \cdot 7\ H_2O$, (**b**) 50 μM TPEN, or (**c**) 40 μM CP. (**a**), (**c**), and (**e**) The data represent the mean from three independent experiments. Error bars have been omitted for clarity. (**b**), (**d**), and (**f**) Quantification of data in (**a**), (**c**), and (**e**), respectively, using area under the curve (AUC). Data are the mean ± SD from three independent experiments. The same lowercase letters indicate samples that are not significantly different and different lowercase letters indicate significant differences ($p$<0.05).

**Table S1** Strains and plasmids used in this study

| Strain or plasmid | Lab Strain | Description | Source |
|---|---|---|---|
| ***P. aeruginosa* strains** | | | |
| PAO1 | DH1856 | Wild type (WT) | (1) |
| PAO1 *att*::P$_{PA3600}$-*lacZ* | | PAO1 (DH1856) expressing P$_{PA3600}$-*lacZ* promoter fusion at the *att*::Tn7 site | This study |
| PAO1V | DH3124 | Wild type (WT), Carb$^{R}$ | (2) |
| PAO1V Δ*lasA* | DH3125 | PAO1V (DH3124) with *lasA* replaced by a gentamicin resistance cassette, Gm$^{R}$ | (2) |
| PAO1V Δ*lasA*+*lasA* | | PAO1V Δ*lasA* complemented with *lasA* on pMQ70_*lasA* | This study |
| PAO1V Δ*lasAB* | DH3126 | PAO1V Δ*lasA* (DH3125) with *lasB* replaced by a streptomycin resistance cassette, Gm$^{R}$, Strep$^{R}$ | (2) |
| ***S. aureus* strains** | | | |
| SH1000 | DH2582 | MSSA 8325-4 with *rsbU* restored | (3) |
| ***E. coli* strains** | | | |
| S17 λpir | DH71 | Used as a conjugation partner for introducing pMQ30- and GH121-based plasmids | |
| DH5α | DH51 | Used for plasmid preparations | |
| NEB 5α | | DH5α derivative used for plasmid preparations | NEB |
| T7 Express | | Used for T7 protein expression | NEB |
| **Plasmids** | | | |
| pMQ30 | DH2620 | Suicide vector for allelic replacement; Gm$^{R}$ | (4) |
| GH121 | DH2830 | Modified pMQ30 (DH2620) with *att* homology for inserting sequences at the *att*::Tn7 site via allelic replacement; Gm$^{R}$ | (5) |
| GH121_P$_{pqsA}$-*lacZ* | DH3785 | *lacZ* under control of the *pqsA* promoter, for integration at the att::Tn7 site; Gm$^{R}$ | (6) |
| GH121_P$_{PA3600}$-*lacZ* | DH3229 | *lacZ* under control of the *PA3600* promoter, for integration at the att::Tn7 site; Gm$^{R}$ | This study |

**Table S1** continued

| | | | |
|---|---|---|---|
| pMQ70 | DH1682 | Vector for arabinose-inducible gene expression; $Amp^R$ | (4) |
| pMQ70_*lasA* | | Vector for arabinose-inducible expression of *lasA*; $Amp^R$ | This study |
| pET21-S100A8-S100A9 | | Vector for IPTG-inducible co-expression of S100A8 and S100A9; $Carb^R$ | (7) |

**Table S2** Primers used in this study

| Primer purpose and name | Sequence (5'-3') |
|---|---|
| **GH121_P$_{PA3600}$-*lacZ* construction** | |
| P$_{PA3600}$_fwd | GCGATTGACGGCGGGCGTCGCGATCGCCGGGGCCGCATGACTGCCTCGGCTTCCAGCAGGTACTG |
| P$_{PA3600}$_rev | TTGGGACAACTCCAGTGAAAAGTTCTTCTCCTTTACTCATGGGGATGCCTCCTACATAATG |
| **pMQ70_*lasA* construction** | |
| pMQ70_fwd | GATCCTCTAGAGTCGACCTG |
| pMQ70_rev | TACCGAGCTCGAATTCGC |
| PA1871_fwd | TGGGCTAGCGAATTCGAGCTCGGTAATGCAGCACAAAAGATCCCGCG |
| PA1871_rev | GCCTGCAGGTCGACTCTAGAGGATCTCAGAGCGCCAGGCCGGG |
| pMQ70_Seq_fwd | ACCTGACGCTTTTTATCGCAAC |
| pMQ70_Seq_rev | TTATCAGACCGCTTCTGCGTT |
